# The Cryo-EM Structures of two Amphibian Antimicrobial Cross-β Amyloid Fibrils

**DOI:** 10.1101/2022.01.08.475498

**Authors:** Robert Bücker, Carolin Seuring, Cornelia Cazey, Katharina Veith, Maria García-Alai, Kay Grünewald, Meytal Landau

## Abstract

The amyloid-antimicrobial link hypothesis is based on antimicrobial properties found in human amyloids involved in neurodegenerative and systemic diseases, along with amyloidal structural properties found in antimicrobial peptides (AMPs) across kingdoms of life. Supporting this hypothesis, we here determined the fibril structure of two AMPs from amphibians, uperin 3.5 and aurein 3.3, by cryogenic electron microscopy (cryo-EM), revealing amyloid cross-β fibrils of mated β-sheets at atomic resolution. Uperin 3.5 displayed substantial polymorphism with a protofilament of two mated β-sheets. The determined structure was a polymorph showing a 3-blade symmetrical propeller of nine peptides per fibril layer including tight β-sheet interfaces. This cross-β cryo-EM structure complements the cross-α fibril conformation previously determined by a crystal structure, substantiating a secondary structure switch mechanism of uperin 3.5. The aurein 3.3 arrangement consisted of six peptides per fibril layer, all showing kinked β-sheets allowing a rounded compactness of the fibril. The kinked β-sheets are similar to LARKS (Low-complexity, Amyloid-like, Reversible, Kinked segments) found in human functional amyloids. The amyloidal properties of antimicrobial peptides shed light on a mechanism of regulation of animicrobial activity involving self-assembly and fibril morphological variations. Moreover, the known endurance of amyloid structures can provide a template for the design of sturdy antimicrobials.

## Introduction

Antimicrobial peptides (AMPs) are a diverse group of molecules that evolved in most organisms to combat microbial challenge since the early days of life. Thousands of different AMPs were already identified in many branches of the tree of life, with very little homology in sequence and in length^1^. Certain AMPs assemble into well-ordered fibrils that resemble amyloids^2–9^, which typically form cross-β fibrils composed of tightly mating β-sheets, often displaying anhydrous interfaces called steric zippers^10^. Amyloids are best known for their association with neurodegenerative and systemic diseases^11^. However, various organisms secrete amyloids that carry diverse physiological roles, including in storage of peptide hormones, formation of membrane-less organelles, and as toxins^12–24^.

Most amyloids, functional and pathological, share a β-rich structure, often with mated sheets yielding different polymorphs of cross-β interfaces. Another form of amyloid, named cross-α, is composed entirely of amphipathic α-helices that stack perpendicular to the fibril axis into mated ‘sheets’, an arrangement reminiscent of the cross-β fibrils. The cross-α form was revealed by the crystal structure of the cytotoxic phenol-soluble modulin α3 (PSMα3) peptide secreted by the pathogenic *Staphylococcus aureus* bacterium^25^, and recently, by the crystal structure of the antibacterial peptide uperin 3.5 secreted by *Uperoleia mjobergii* (Mjoberg’s toadlet), which forms fibrils with amyloid properties^7,26–28^. The cross-α amyloid was suggested to serve as a toxic functional amyloid form^7,29,30^. Biophysical studies, including secondary structure analysis using solid-state circular dichroism, Fourier-transform infrared spectroscopy, and fiber X-ray diffraction, showed a secondary structure switch between α-helical and β-rich fibrils depending on environmental conditions, in particular the presence of lipids which drive a transition to an α-rich state^7^. Here, a cryo-EM study of uperin 3.5 in the absence of lipids reveals the presence of β-rich fibril polymorphs, and the atomic details of the cross-β configuration, allowing for a direct comparison to the cross-α fibril and exposing the substantial polymorphism of the same sequence.

A recently developed computational screen aimed to identify fibril-forming AMPs (ffAMPs) additional to uperin 3.5 found fifteen new sequences, including aurein 3.3, another amphibian AMP secreted by *Ranoidea raniformis* (Southern bell frog) (Ragonis-Bachar, Rayan *et al, In preparation*). Here we show the cryo-EM structure of aurein 3.3, revealing a cross-β fibril with kinked β-sheets. Such sharp kinks in the peptide backbone are the hallbark of Low-complexity Amyloid-like Reversible Kinked Segments (LARKS) within functional amyloids^16–20,31^.

The ability of uperin 3.5, aurein 3.3, and other AMPs secreted by different organisms, to assemble into amyloid-like fibrils^2–9^, complement the observation that human amyloids associated with neurodegenerative and systemic diseases, including Amyloid-β, tau, and α-synuclein associated with Alzheimer’s and Parkinson’s diseases, respectively, possess antimicrobial properties^32–42^. Here, the cryo-EM structures of uperin 3.5 and aurein 3.3 provide an atomic-level validation for the formation of cross-β amyloid structures by AMPs, supporting the amyloid-antimicrobial link. Moreover, the structures may direct the development of sturdy antimicrobials with high stability and adherence to surfaces.

## Results

For both of the studied ffAMPs, cryo-EM data unambiguously prove an amyloid cross-structure comprised of -sheets with tightly coupled dry interfaces and strands perpendicular to the fibril axis. While for both ffAMPs pronounced fibril polymorphism is observed, the power spectra of all significantly populated two-dimensional class averages exhibit a pronounced peak near 4.85 Å, indicating that all observed polymorphs are of a canonical cross-structure. For each ffAMP, we identified the most abundant polymorph, and solved its three-dimensional structure through helical single-particle analysis.

### Cryo-EM structure of uperin 3.5

Using two-dimensional class averaging of fibril segments extracted from the electron micrographs, we identified a number of polymorphs with markedly different appearances (Figure 1). While the unique morphology of the most abundant polymorph (type I; relative abundance 71%) makes both individual fibrils in the micrographs and two-dimensional class averages easy to identify, variations between the other fibril types are more subtle. To identify those, we applied the CHEP algorithm^43^ to the two-dimensional classification of segments, as well as three-dimensional classification in later analysis steps. From those steps, we distinguished additional three clearly identifiable minority polymorphs (fibril types IIa, IIb, IIc; relative abundance 5%, 15%, 9%, respectively). Two-dimensional class averages from segments of fibrils from each type, along with lateral cuts through each reconstructed three-dimensional volume are shown in Figure 1a-d. We found those polymorphs to be structurally related in that they are assembled from a protofilament (type IIa) comprising two symmetric -sheets, which are mated near their termini. Neither between the sub-types of type II, nor between types I and II did we find a transition of one polymorph to another within the same fibril (Figure 1f). For all helix types, layer lines normal to the filament axis can be observed at a spacing of approximately 4.85 Å, as is evident from the power spectra of class averages along the helical axis in Figure 1e. This type of stacking is a hallmark of cross-fibrils, suggesting an amyloid-like structure of all observed uperin 3.5 polymorphs.

**Figure 1:**
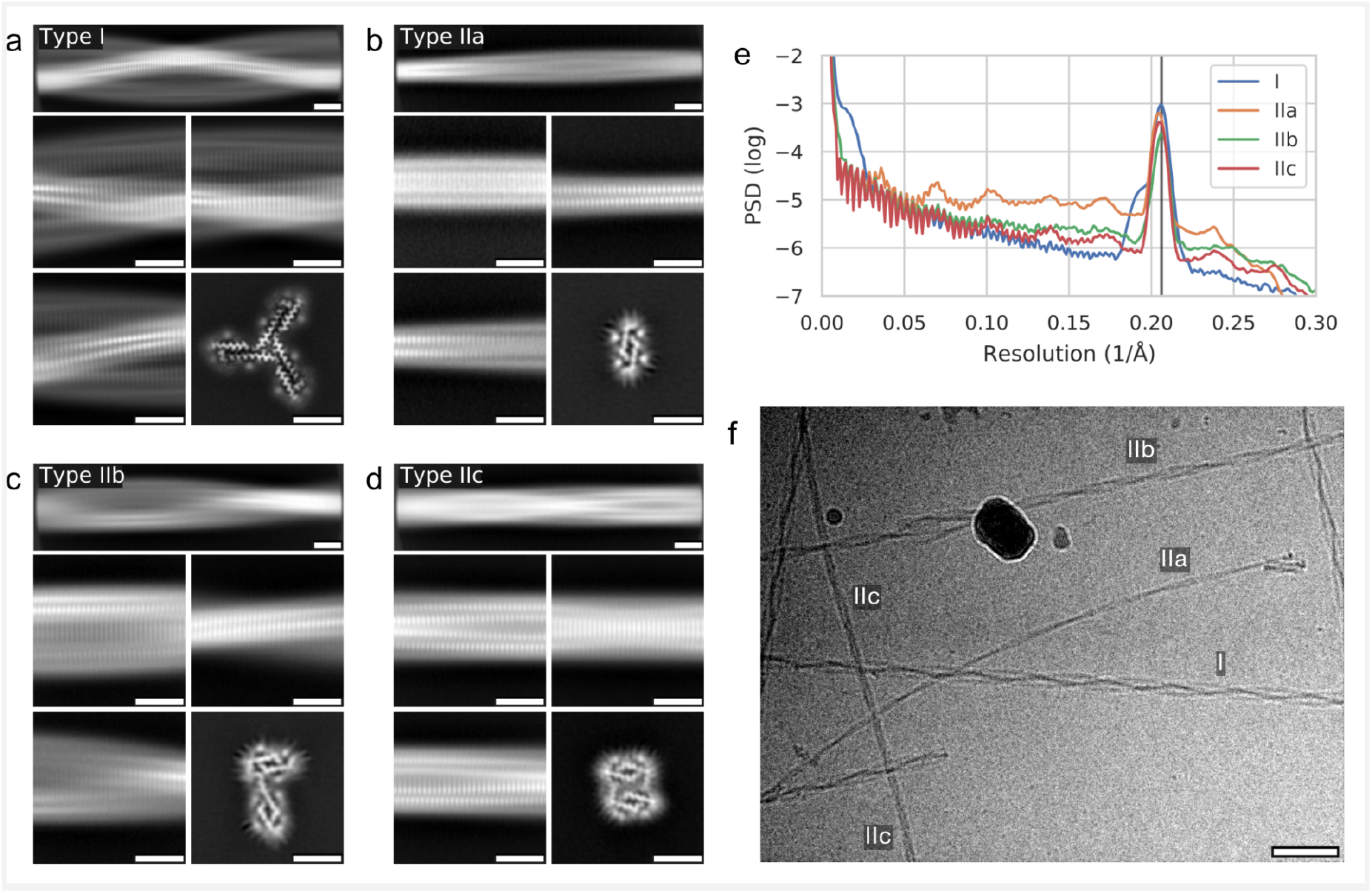
Polymorphs of uperin 3.5 fibrils. (a) Representative two-dimensional segment averages over long (top row) and short (middle row, bottom left) ranges from fibril types I, and lateral cut through three-dimensional reconstructed volume (bottom right). Scale bars are 5 nm. (b-d) As (a), but for fibril types IIa, IIb, and IIc, respectively. (e) Power spectra of two-dimensional class averages taken along the fibril axis, shown on a logarithmic scale. The vertical line corresponds to the helical rise of 4.85 Å. (f) Representative cryo-EM micrograph containing specimens of all types. Scale bar is 50 nm.

For the uperin-3.5 helix-type II polymorphs, despite the presence of a pronounced peak near the -sheet inter-strand spacing of about 4.85 Å in 2D class averages, we were not able to improve the resolution of 3D reconstructions sufficiently to determine the staggering structure of strands. Still, the cross sections of the initial models as shown in Figure 1 are compatible with the sequence of the peptide monomer and give valuable insight into the lateral structure of the fibrils.

We determined the three-dimensional structure of type I fibrils using helical single-particle analysis to high resolution as described in the online methods. The reconstructed volume at an average resolution of 2.97 Å is shown in Figure 2. We find a propeller-shaped structure with threefold rotational symmetry about the fibril axis, where each of the three asymmetric units per filament layer comprises three chains. An atomic model of the uperin 3.5 peptide built into each asymmetric unit (propeller blades) shows excellent agreement with the data in the inner section of the fibrils, comprising full-length (17 residues) monomers in chain A, and 14 and 7 residues in chains B and C, respectively (chains represent the layers along each -sheet, named as indicated in Figure 2). As seen in Figure 3, chains A and B are tightly interdigitated and form -sheets along the fibril axis, both hallmark features of a canonical amyloid cross-fibril. Chain C is almost perpendicular to the other two. In the outer regions of chains B and C, the observed density becomes increasingly blurry, suggesting flexible terminal sections, which we did not include in the model.

**Figure 2:**
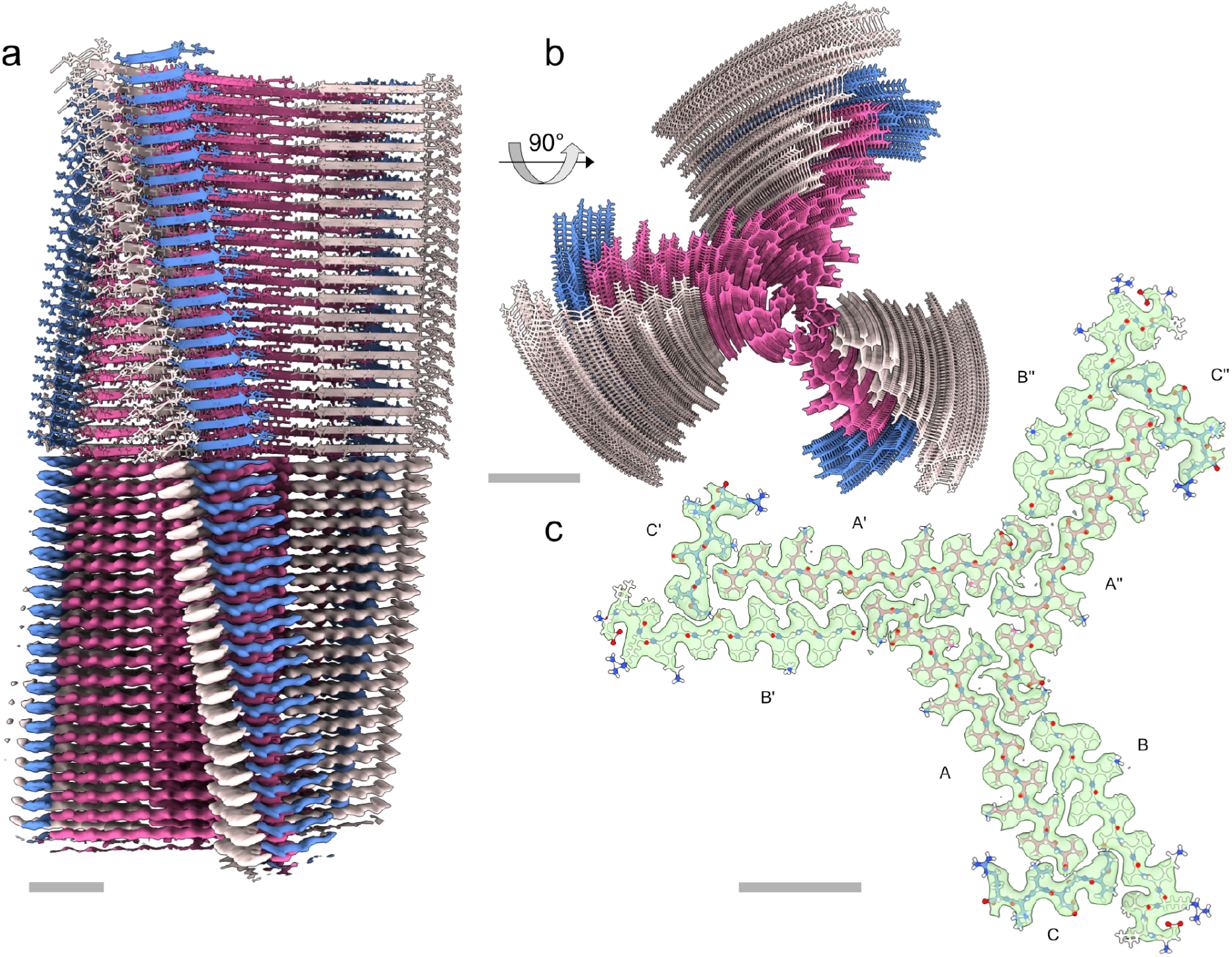
Cryo-EM structure of uperin 3.5 type I. (a) Reconstructed coulomb density (lower half) and cartoon representation (upper half) of 40 fibril layers. Coloring is by peptide monomer, taking into account the helical and rotational C3 symmetries. (b) View along the fibril axis. (c) Atomic model of a single layer built into the coulomb density with chain identifiers as discussed in the text. Scale bars are 20 Å.

**Figure 3:**
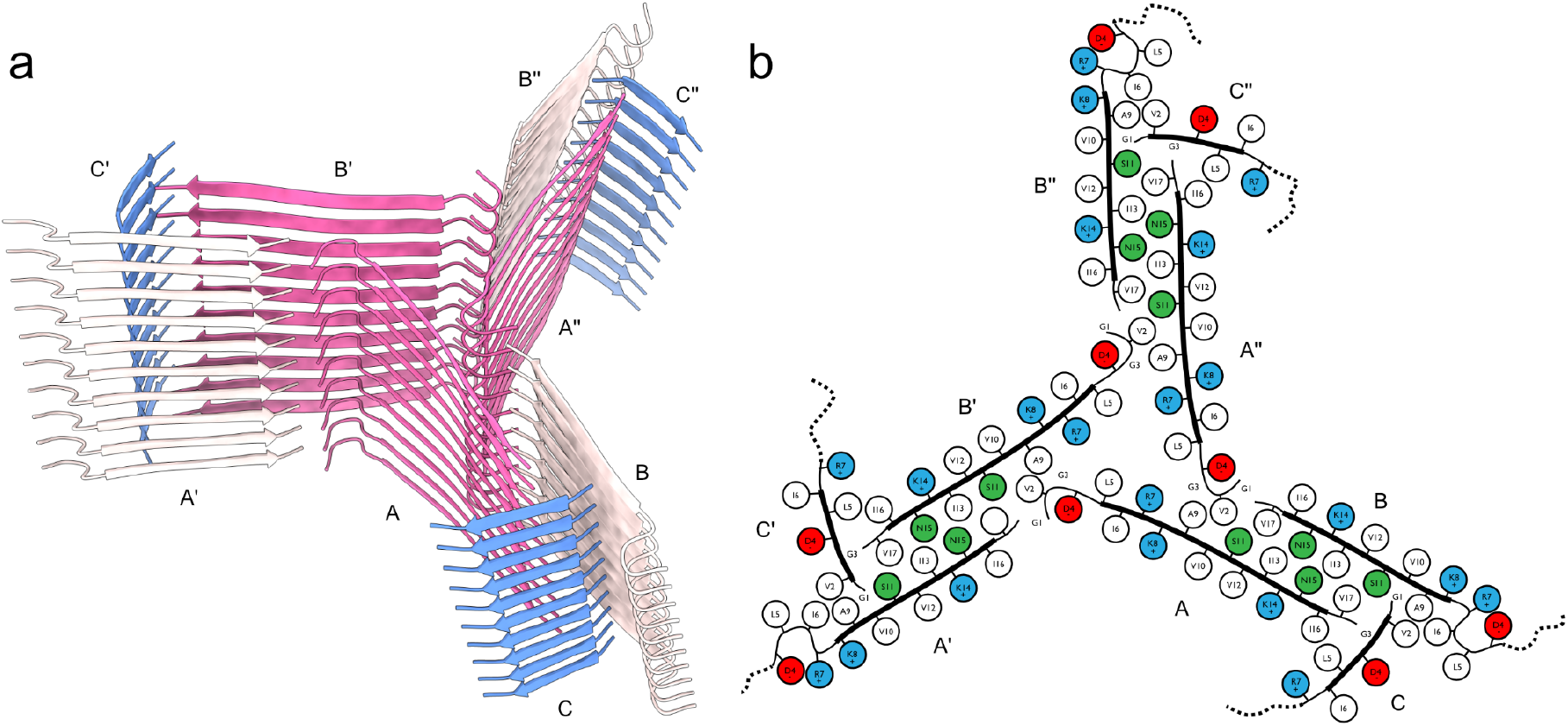
Fibril structure of uperin-3.5. (a) 10-layer section of the cross-beta structure shown as beta-sheet cartoons at an inclination of 40° with respect to the fibril axis. (b) Cartoon of residue properties within the fibril cross-section. Hydrophobic, polar, and negatively/positively charged residues are indicated in white, green, red, and blue, respectively. Dotted lines represent unmodeled flexible terminal sections.

Since amyloid fibrils are often unusual in that pairs of β-sheets mate more closely than the adjoining surfaces in other protein complexes, quantitative measures of amyloid stability are based on solvent-accessible surface area (SASA) buried at the interface between the mating sheets. The overall packing of the uperin 3.5 fibril is highly dense, with the SASA buried for a single layer (fibril cross-section of nine lateral chains) covering 13570 Å^2^, which is 82% of the total area of this layer. The inner chains (A, A’, and A’’) show the largest SASA buried by surrounding chains, as expected by their mid-fibril location (Table 1;Figures 2&3). On average, 69% of the total area of individual chains (A, B and C) in the fibril is buried by surrounding chains. The average SASA for an individual chain buried by all axially neighboring chains (from the same β-sheet), or all laterally neighboring chains (all chains excluding those from the same β-sheet) constitute 59% and 26% of the area of an individual chain, respectively (Table 1). This indicates that most area is buried along the fibril axis compared to lateral associations, which is expected considering that each chain is buried by top and bottom axial interactions along the fibril. Considering that only 14 and 7 residues, out of 17-residues, are visible for chains B and C, respectively, the indicated numbers in Table 1 do not represent the actual area buried in the full-length peptides. The percentage of area buried in the lateral associations will probably be effectively smaller. However, as the axial associations along the sheet are the most contributing factor, and should be similar along the entire chain, we do not expect the overall percentage of buried SASA to change significantly.

**Table 1:**
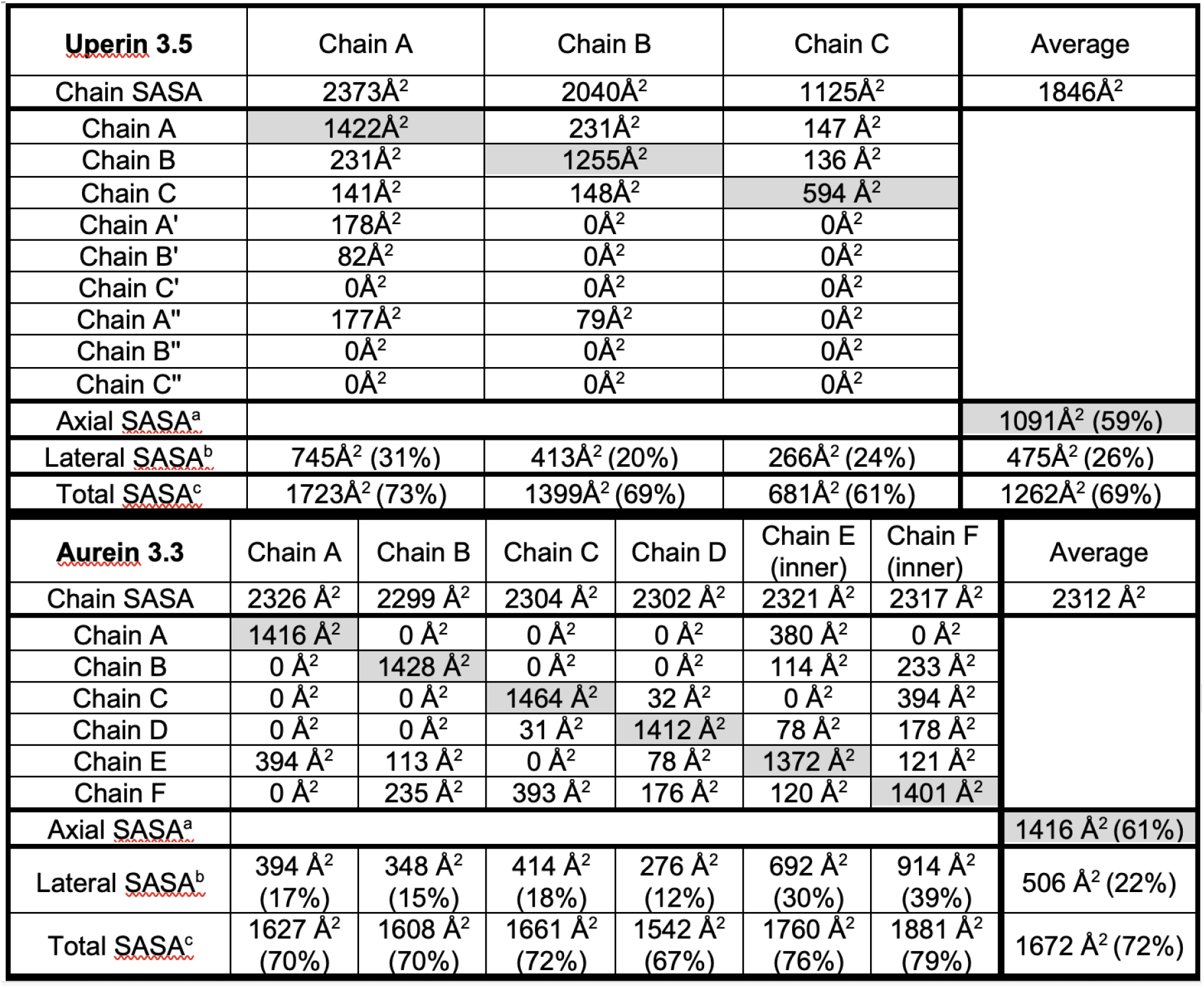
solvent-accessible surface area (SASA) buried within the uperin 3.5 and aurein 3.3 fibrils.

The center of the propeller is composed of inward-facing Arg7 and Leu5 residues from chains A, A’ and A’’, creating a positively charged and hydrophobic core (Figure 3). The C-terminus of uperin 3.5 was synthesized in the amidated form, according to the natural presence in uperin 3.5. This amide group can form inter-strand hydrogen bonds between the amine and carboxyl along the fibril axis. The amine in the C-terminus of chain B can form a potential hydrogen bond with the backbone oxygen of Gly1 of chain A from an adjacent blade, overall connecting these termini located in the core of the fibril. The amine in the C-terminus of chain A can form a potential hydrogen bond with the backbone oxygen of Gly3 of chain C. The interface between the tightly mated β-sheets of chains A and B in each propeller blade is composed of mostly hydrophobic residues, except from Asn15 forming inter-sheet hydrogen bonds, and Ser11. In chain A, Ser11 is facing the hydrophobic interface, while in chain B, Ser11 forms potential hydrogen bonds with the N-terminus of chain C. The N-terminal part of chains A and B is located outside the mated interface. Asp4, located in this external region, is forming a potential hydrogen bond or a salt bridge with Arg7 within chain B, yielding a bent structure. The first 3-residues of chain B, probably extending the fibril surface with a solvent exposed N-terminus, are not observed in the map. In the more extended chain A, the N-terminus forms a potential salt bridge with Asp4, satisfying charges in the center of the fibril. In chain C, Asp4 is protruding towards the surface of the fibril, contributing a negative charge to an otherwise positively charged and hydrophobic fibril surface. Overall, the inter-sheet interfaces within the fibril are mostly hydrophobic but also contain polar interactions. The outer surface and fibril inner circular core are composed of mostly hydrophobic and positively charged patches (Figure 3).

### Cryo-EM structure of aurein 3.3

In contrast to the case of uperin 3.5, we did not observe any pronounced polymorphism of fibrils formed by the antimicrobial peptide aurein 3.3. Following similar sample preparation, data collection and processing steps as for uperin 3.5, we arrived at the filament structure as shown in Figure 4.

**Figure 4:**
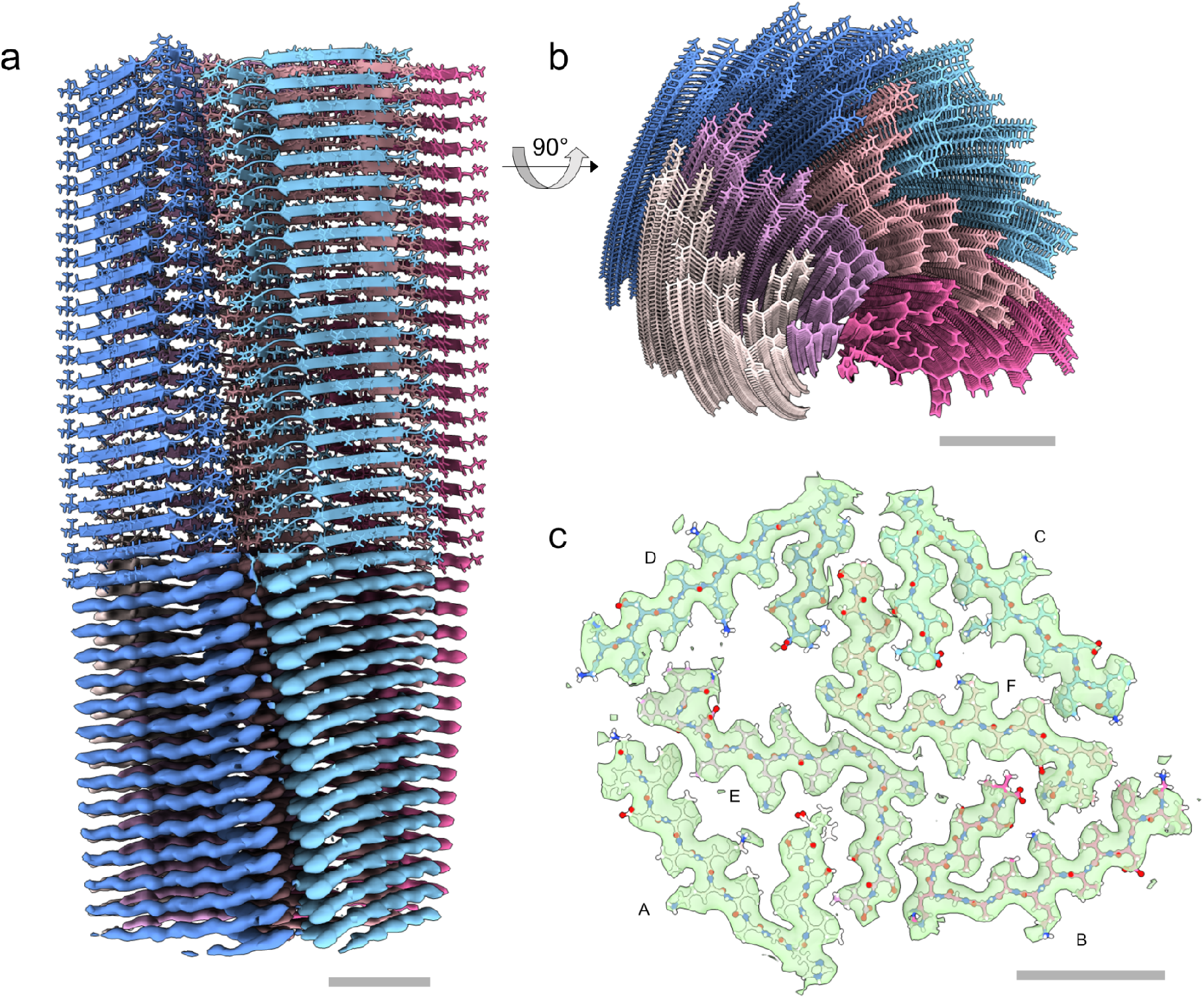
Cryo-EM structure of aurein 3.3. (a) Reconstructed coulomb density (lower half) and cartoon representation (upper half) of 40 fibril layers. (b) View along the fibril axis. (c) Atomic model of a single layer built into the coulomb density with chain identifiers as discussed in the text. Scale bars are 20 Å.

Unlike the cross-structure with extended mated regions found in uperin 3.5 type I filaments, aurein 3.3 fibrils exhibits a complex, unusual in-plane structure with six chains per helical layer, each corresponding to a single full-length 17-residue peptide, as highlighted in Figure 5. Two of those (chains E, F) are arranged in a cross-like shape conforming to C2_1_ screw-symmetry along the fibril axis and form kinked -sheets in their inner section which closely meet at a roughly 90° bend near Gly11. Wrapped around this inner cross are four more strands of mutually similar conformation, each having a sharp in-plane kink near the outward-facing His12 residue, which is enabled by Gly11 leaving a void into which Val14 can fold. The outer chains cross the fibril normal at an unusually large angle, which, together with the screw symmetry of the inner cross, leads to a complex staggered structure of the fibril without well-defined rungs. The symmetry of the inner cross is broken due to the stark dissimilarity of the orientation of the outer chain B compared to the other outer chains A, C and D (chains represent the layers along each -sheet, named as indicated in Figure 5).

**Figure 5:**
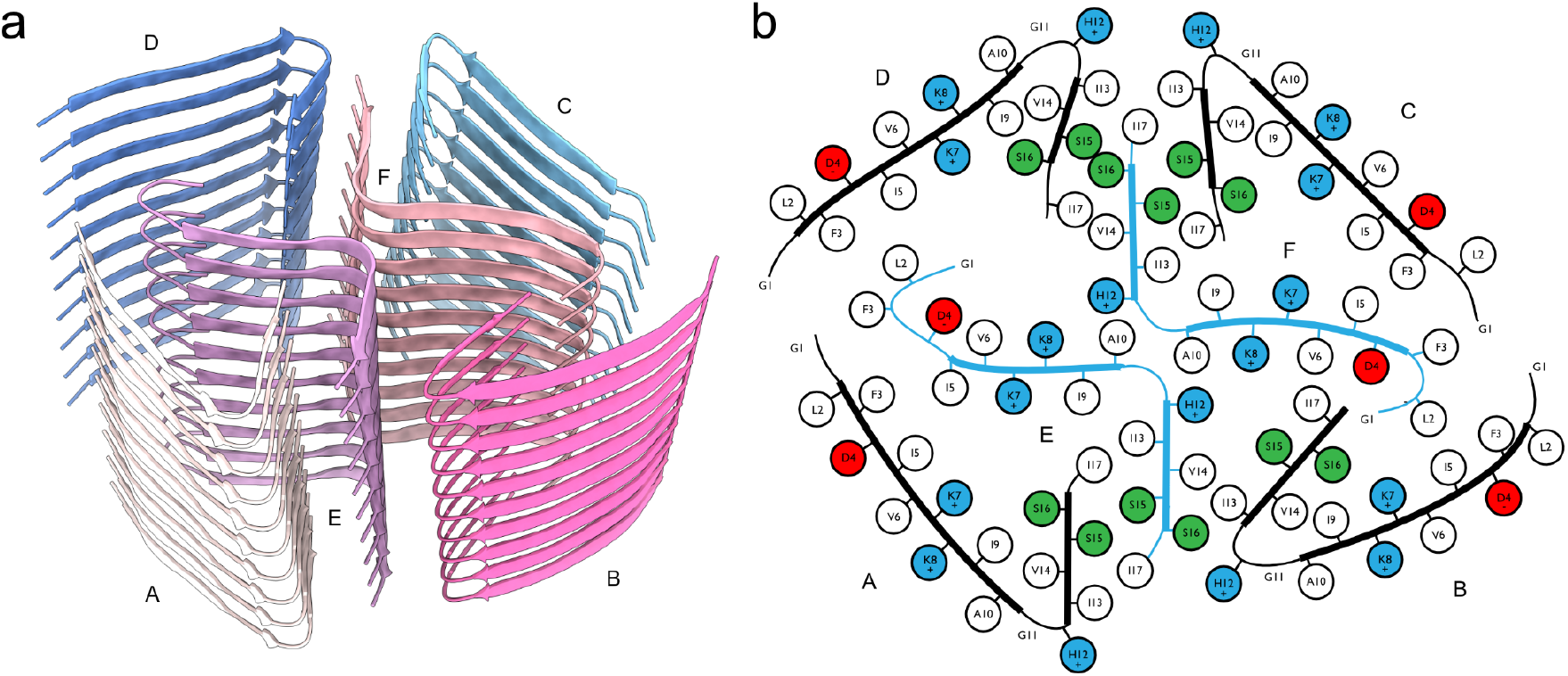
Fibril structure of aurein-3.3. (a) 10-layer section of the cross-beta structure shown as beta-sheet cartoons at an inclination of 40° with respect to the fibril axis. (b) Cartoon of residue properties within the fibril cross-section. Hydrophobic, polar, and negatively/positively charged residues are indicated in white, green, red, and blue, respectively. Note that due to the strongly staggered arrangement of the beta-sheet strands along the fibril, distances between residues from different sheets may appear smaller than they are.

**Figure 6:**
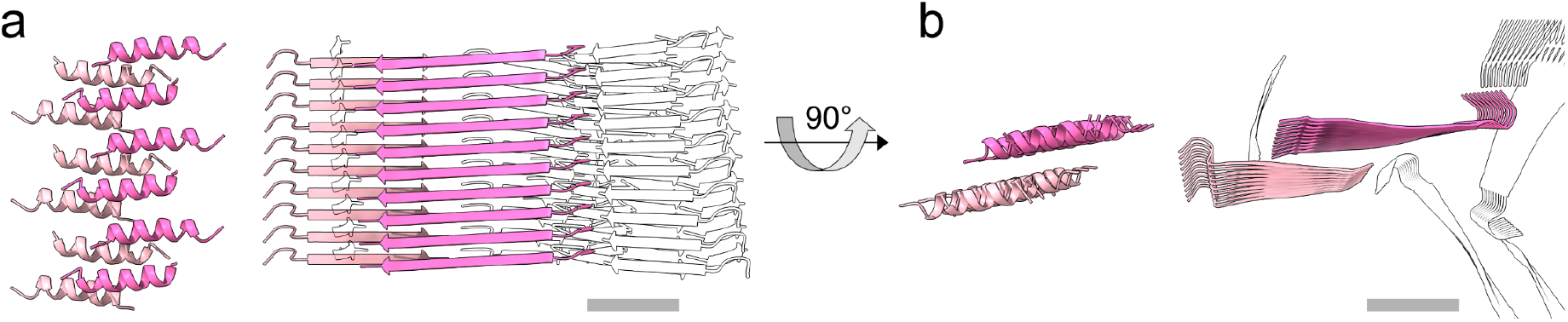
Uperin 3.5 cross-α and cross-β structures. Side view (a) and top view (b), showing a 10-layer section of the crystallographic cross-α (left) and cryo-EM cross-β (right) fibril structures, respectively. The crystal structure of uperin 3.5 exposed cross-α fibrils composed of α-helices arranged in ‘anti-parallel sheets’ that mate via a hydrophobic interface (PDB ID 6GS3)^2–7^. For the cross-β structure, two of the closely mated sheets (chains A and B) of a type I helix are highlighted; the others are shown as silhouettes. All models to scale; scale bars are 20 Å.

The overall packing of the aurein 3.3 fibril is highly dense, with the SASA buried for a single layer (fibril cross-section of six lateral chains) covering 9802 Å^2^, which is 83% of the total area of this layer. The two inner chains (E, F) show the largest SASA buried by surrounding chains, as expected by their mid-fibril location (Table S1,Figure 4, Figure 5). The average SASA for an individual chain buried by all surrounding chains constitutes 73% of the total area of the chain. The average SASA for an individual chain buried by all axially neighboring chains (from the same β-sheet), or all laterally neighboring chains (all chains excluding those from the same β-sheet) constitute 63% and 21% of the area of the chain, respectively (Table S1). This indicates that most area is buried along the fibril axis compared to lateral associations. Nevertheless, the lateral contacts among the six -sheets (fibril cross-section) are rather extensive, with the two inner chains showing relatively high percentage of buried area (Table S1).

In the outer chains C and D, the C-terminal part of the kinked -sheet forms tightly mated -sheet interactions with the C-terminal part of the inner chain F, overall showing a stable interface of three short -sheets, resembling lateral association between -sheets in the crystal structures of short amyloid segments^44,45^. Similarly, the C-terminal part of the kinked -sheet of chain A forms tightly mated -sheet interactions with the C-terminal part of the inner chain E. Chain B, in contrast to the other outer chains, more tightly fills the void spanned by both inner chains, E and F, contacting both (Table S1), yet with no mated -sheet interface. Overall, the SASA buried for the outer chains is similar, with chain D showing the smallest contact area, being less tightly packed compared to the other chains. Of note, Phe3 from chains A, D and E are tightly packed, and similarly, also Phe from chains B, C and F, contributing to the extensive hydrophobic/aromatic interactions stabilizing and structure. Overall, despite the lack of extended internal mated surfaces, all chains meet near a large number of vertices as seen from Figure 5, which might explain a sufficient degree of stability causing the absence of any other polymorphs in our sample.

The aurein 3.3 fibril contains numerous hydrogen bonds, mostly involving N-O backbone atoms between the -strands along the -sheets. Side chain interactions involve hydrogen bonds between Ser15 from chain D and Ser16 from chain F, and between Ser15 pairs from chains C and F. In the two inner chains (E and F), the N-terminus is bent, such that Asp4 can form a potential salt bridge with the N-terminal charged amine. In the outer chains, the N-terminal part is more extended and Asp4 is protruding outwards. Lysine and histidine residues are facing both the inner and outer parts of the fibril. Between the two inner chains, Lys8 and His12 are facing into the core of the fibril. Lys7 is facing the direction of another Lys7 from the outer chains A and C. The relative lateral closeness of lysine residues, in addition to their stacking along the fibril axis, might suggest an uncharged form, even around neutral pH, as suggested for other amyloid structures with lysine ladders^46^. Lys8 and His12 from the outer chains protrude outwards from the surface of the fibril. Overall, the lateral association of -sheets in the fibril shows very few potential polar interactions and is mostly stabilized by van der Waals forces.

The solvent-accessible surface area (SASA) buried per chain was calculated at its interface with different sets of other chains as defined: a) The “Axial SASA” refers to the SASA buried by surrounding chains on the same sheet, indicated with the gray shading between all pairs. b) “Lateral SASA” refers to the SASA buried per chain by all other surrounding chains except the ones from the same -sheet. Percentage of this buried area from the total SASA of the chain (first row) is indicated. a) “Total SASA” refers to the SASA buried per chain by all other surrounding chains in the fibril. The Percentage of this buried area from the total area of the chain (first row) is indicated.

## Discussion

### Sequence properties and interfaces of the two amphibian AMPs compared to pathological and functional amyloids

The collection of high resolution structures of pathological and functional human amyloids determined thus far (reviewed in^47,48^) support a common cross-β motif, but with differences in the polarity and flexibility of the inter-fibril interfaces, which subsequently impact fibril reversibility and regulation^47^. Despite the dissimilarity in lateral fibril structure and the difference of extended versus kinked β-sheets, the 17-residue ffAMPs uperin 3.5 and aurein 3.3 are remarkably similar in their residue composition. Both contain nine hydrophobic residues, two glycines, one negatively charged aspartate, three positively charged residues (lysine, arginine, or histidine), and two polar residues (serine or asparagine). Compared to a selection of pathological and functional amyloids^47^, the percentage of apolar and of positively charged residues in the two amphibian ffAMPs is higher, while the percentage of negatively charged and polar residues is lower. The amphibian ffAMPs show higher and lower percentage of glycines compared to a selection of pathological and functional amyloids, respectively^47^, which might indicate the intermediate flexibility of the interfaces of the amyloidal ffAMPs. Overall, compared to other amyloids^47^, the sequence content of the two AMPs indicates a dominant apolar and positively charged nature, which fits the functional interactions with negatively charged bacterial membranes.

### Comparison of cross-β cryo-EM structure to the cross-α crystal structure of uperin 3.5

Uperin 3.5 is a rare example of secondary structure polymorphism in the fibril form, with a switch induced by environmental conditions, particularly the presence of lipids^7^. It is also a rare example of the ability of two different structureal biology methods, X-ray crystallography and cryo-EM, to expose two fibril polymorphs principally different in their secondary structure. Biophysical studies previously suggested that uperin 3.5, in the absence of lipids, assumes a cross-β configuration^7^. The atomic details of one of the cross-β polymorphs, and the existence of other β-rich polymorphs, were revealed here (Figures 1-3). In contrast to the cryo-EM structure reported here, in which uperin 3.5 was dissolved in water, crystallization conditions^7^ included a polyether based on polypropylene glycol, which might have stabilized the amphipathic cross-α conformation. Interestingly, the structurally similar polyglycerols were suggested to serve as building blocks for amphiphilic structures and to affect fibrillation kinetics of amyloid fibrils^49^. Overall, crystallization most likely favored a conformation more prone to forming crystals, such as the cross-α configuration with tight packing of the α-helical sheets, compatible with ordered crystal packing. This finding highlights the importance of single-particle studies in unravelling structural polymorphism of amyloid filaments and supramolecular assemblies.

### Uperin 3.5 fibril structure and antibacterial activity

Uperin 3.5 is secreted on the skin of toadlets, and it is likely that secreted monomers having disordered structures that then aggregate into a stable form^7^. It is still unclear to what level the toxicity differs between cross-β and cross-α uperin 3.5 fibril configurations. The presence of bacterial cells might induce fibrillation that enhances toxicity against the bacterial cells and eliminates the threat of infection^7^. A model in which the cross-β configuration is less toxic is reminiscent of human amyloids for which the mature β-rich fibrils are considered non-toxic whereas toxicity is attributed to prefibrillar oligomeric conformations^50^, which might contain α-helices^51,52^. However, the cryo-EM structure of uperin 3.5 displays positively charged and hydrophobic patches composing the surface of the fibril (Figure 3), making them prone for interactions with bacterial membranes, as proposed for other, non-amyloid, antimicrobial peptide fibrils^53^. It is also possible that different polymorphs of the cross-β type display different levels of toxicity, as human amyloid polymorphism has been correlated with different toxicity levels and prion disease strains^54,55^. Overall, determinants of activity are probably complex, dictated by the presence of specific lipid compositions, which can affect fibrillation rate and polymorphism in the fibrils, including induction of secondary structure switches. Single-particle structures of uperin polymorphs upon interaction with various lipids will be the subject of future studies.

### Reversible interfaces of functional amyloids and kinked β-sheets of aurein 3.3

The structure of aurein 3.3 demonstrated exceptionally sharp kinks in its β-sheets (Figure 4). Kinked β-sheets were observed in short amyloid segments from low-complexity domains of RNA-binding proteins which serve as functional amyloids forming membrane-less organelles^16–20,31^. The kinked β-sheet structural motif, termed LARKS, was also recently observed by a crystal structure of a segment located within the low complexity threonine-rich domain of the *Candida albicans* Als5 adhesin^56^. LARKS are stable, but likely not to the extent of steric zippers of typical amyloids that form irreversible structures^16–20,31^, allowing labile and reversible fibril formation, to enable a wide range of functionality in various physiological contexts^57^. Overall, accumulative structures of kinked β-sheets in different amyloid-producing organisms, from yeast, to amphibian and human, provide support for the connection between those types of interfaces and functional amyloid activities.

## Conclusion

Overall, the cryo-EM structures of uperin 3.5 and aurein 3.3 validate the formation of cross-β amyloid structures by AMPs and thereby support the link between amyloids and antimicrobial activity. This connection suggests a physiological role in neuroimmunity for amyloids, otherwise known as merely pathological^42^. Moreover, AMPs that form highly stable fibrils as amyloids can open paths to develop antimicrobials with enhanced oral bioavailability, stability under harsh conditions, adherences to surfaces, and longer shelf-life.

## Online Methods

### Sample Preparation

Aurein 3.3 from *Ranoidea raniformis* (Southern bell frog) (Uniprot ID P82396 AUR33_RANRN; sequence GLFDIVKKIAGHIVSSI) and uperin 3.5 from *Uperoleia mjobergii* (Mjoberg’s toadlet) (UniProt ID P82042|UPE35_UPEMJ; sequence GVGDLIRKAVSVIKNIV-NH2) were purchased from GL Biochem (Shanghai) Ltd. as lyophilized peptides, at >98% purity. The peptides were incubated in double distilled water (ddH2O) at a concentration of 5 mg/ml in 1.5-ml Eppendorf tubes at 20° C. The optimal incubation time to produce ordered and dispersed fibers was monitored over time by negative-stain transmission electron microscopy of small aliquots, and determined to be 5 days. This time point was used for the preparation of cryo-samples.

### Electron cryo-microscopy

For cryo-microscopy, 3.5 μl of fiber suspensions (sample concentration 10mg/ml peptide) were deposited on glow-discharged holey carbon grids (Quantifoil R2/1, 300 mesh) at 100% humidity and 4° C, blotted after 15 s of wait time with empirically optimized parameters, and subsequently plunge-frozen in liquid ethane-propane using an FEI Vitrobot Mark IV. Samples were imaged on a Titan Krios G3i (ThermoFisher Scientific) transmission electron microscope, operated at 300 kV and equipped with a Gatan Bioquantum energy filter operated in zero-loss mode (20 eV energy slit width). Images were acquired on a Gatan K3 electron counting direct detection camera (Gatan Inc.) in dose fractionation mode using EPU software at a nominal magnification of 105000x (physical pixel size 0.85 A/px). Data collection details are described in Supplementary Table 1.

### Image processing and polymorph separation

For each of the two samples, motion correction and CTF estimation of the motion-corrected micrographs was performed using the CTFFIND4 implementation in Relion 3.1.3^58^. A set of approximately 400 fibril sections was then manually picked in Relion, and overlapping segment boxes of 328^2^ pixels were extracted and exported for neural network training in crYOLO 1.8-beta^59^. Automatic segment picking with inter-segment distance of 30 pixels and filament tracing was then performed in crYOLO, and found segment coordinates were exported in EMAN2 segmented helix format. Using a custom python script used through the External job type in Relion, those coordinates were used to interpolate the segment coordinates to arbitrary segment spacings for the following extraction steps. Segments were extracted with a box size of 632^2^ pixels, downsampled to 256^2^ pixels (effective pixel size 2.1 Å), at an inter-box distance of 58 Å. Using Relion 3.1.3, 2D class averages were computed as a first step of polymorph identification. For the case of uperin-3.5, segments of type-I helices were clearly standing out in the class averages and could be separated manually. The various sub-types of type-II helices could not as easily be distinguished; hence, the classification results were fed into the CHEP algorithm^43^, which, in brief, clusters the distribution of 2D classes within the segments of each helix, in order to associate each full helix with a putative polymorph type. Using this method, we could distinguish helices of four different sub-types, each of which were subjected to another round of 2D class averaging and segment selection. For the case of aurin-3.3, the additional CHEP step was not required, as only a single polymorph was apparent in the initial 2D classes.

### Initial model building and map refinement

For each helix type separately, segments were then imported into cryoSPARC 3.2^60^ for initial model building using stochastic gradient descent optimization in helical mode, where the helical rise was estimated using the observed helix cross-over distance in the 2D class averages, assuming a helical pitch of 4.8 Å and a left-handed helix.

Next, another set of segments of 328^2^ pixels box size at an inter-box distance of 29 Å were extracted without down-sampling. Starting from the previously determined initial model, all following helical refinement and classification steps were performed in Relion 3.1.3^61^ using this set of segment images.

For uperin 3.5 type I, a sequence of 3D classification, refinement and CTF refinement steps, reducing the number of unique helical units from 9789852 after picking to 860080 led to a final resolution (FSC=0.143) of 2.97 Å (Extended Data Figure 7a). A slight difference in local resolution between the tightly bound inner surfaces of each strand and the fibril exterior is observed (Extended Data Figure 7b). In all refinement and classification steps, tight priors from filament picking on the in-plane (psi) and tilt (rho) angles were imposed. The final volume used for model building was obtained using automatic B-factor sharpening in Relion, providing a soft mask around a 140 Å long section of the helix.

For aurein 3.3, a dihedral D2 pseudosymmetry is present due to the almost right-angle kink in the inner sheets (E, F) and the near-symmetry in the lateral plane of the outer sheets (A-D). This complicates the disambiguation of each filament’s polarity, that is, a 180° flipping of in-plane angles, which did not converge by the end of initial refinement runs, leading to unsatisfactory resolution and unreasonable features on especially the outer peptide strands. We succeeded to disambiguate the polarities by inserting a 3D classification stage, where on top of the in-plane and tilt angles, we imposed a soft prior on the rotation angle along the fibril axis. This led to two equally populated classes at high resolution, one of which similar to previous refinements, and one of which exhibiting a biologically unreasonable structure and discontinuties of rotation angle along the fibrils. After flipping the in-plane angle of particles from the latter, another refinement run yielded converging helix polarities, continuous rotation angles and much improved appearance. To further corroborate the unusual breaking of twofold symmetry along the helical axis by the outer sheets B and D, we added a refinement run into the pipeline, where we modified the initial model derived from the previous run such that the density of sheet D was deleted and replaced by a C2_1_ symmetry-equivalent version of sheet B. We found that despite this edit, the original, asymmetric structure was restored within the first few refinement iterations. After final runs of CTF refinement and Bayesian polishing we arrived at a resolution (FSC=0.143) of 3.48 Å from 141156 unique helical units (Extended Data Figure 8a). Computing local resolution using a sliding-window FSC in Relion revealed the highest resolution to be present near the kinks of the inner -sheet at the center of the fibril, with significantly worse values near the outward-facing residues (Extended Data Figure 8b).

As for uperin-3.5, the final volume used for model building was obtained using automatic B-factor sharpening in Relion, providing a soft mask around a 140 Å long section of the helix.

**Extended Data Figure 7:**
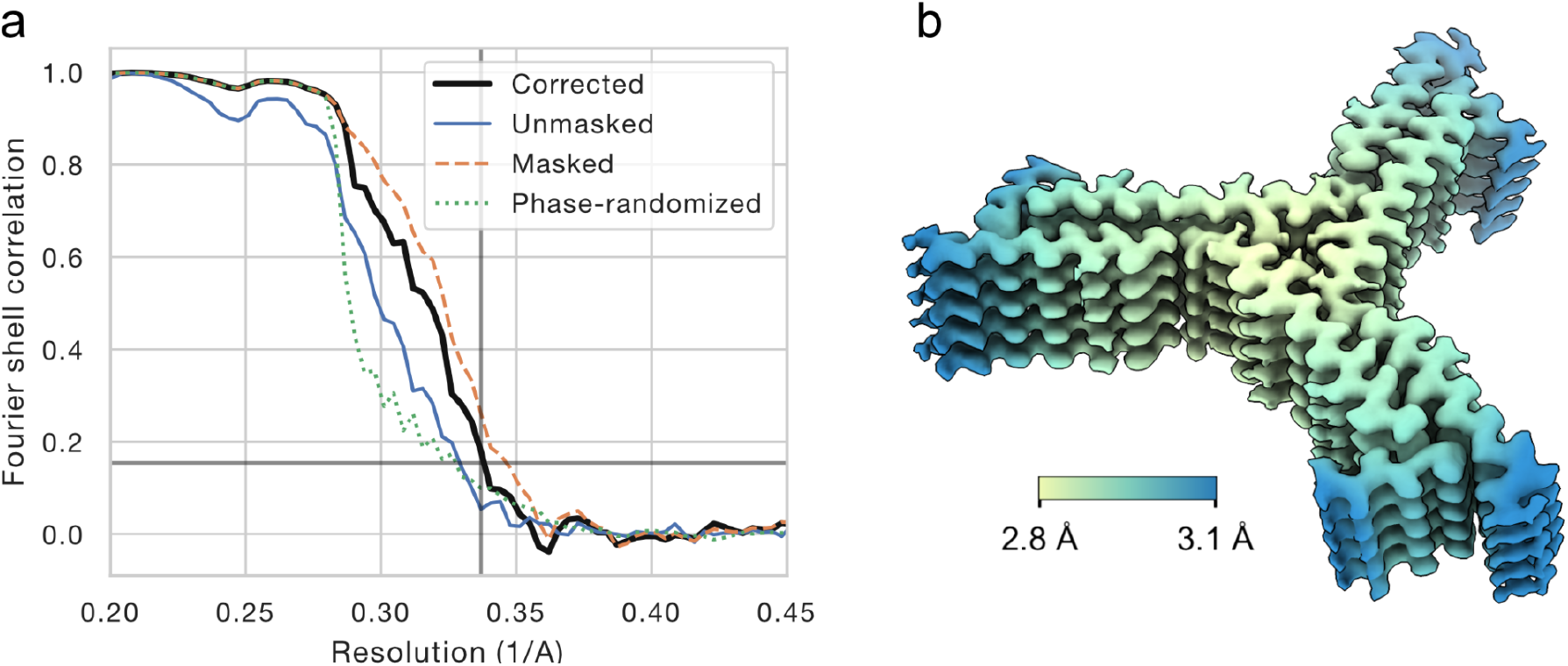
(a) Half-set Fourier shell correlation for uperin 3.5 type I. The grey vertical line indicates the FSC=0.143 resolution of 2.97 Å, for a masked FSC after subtraction of a phase-randomized masked FSC (corrected, black line). (b) Local resolution map, calculated using half-set FSCs within a sliding window in Relion.

**Extended Data Figure 8:**
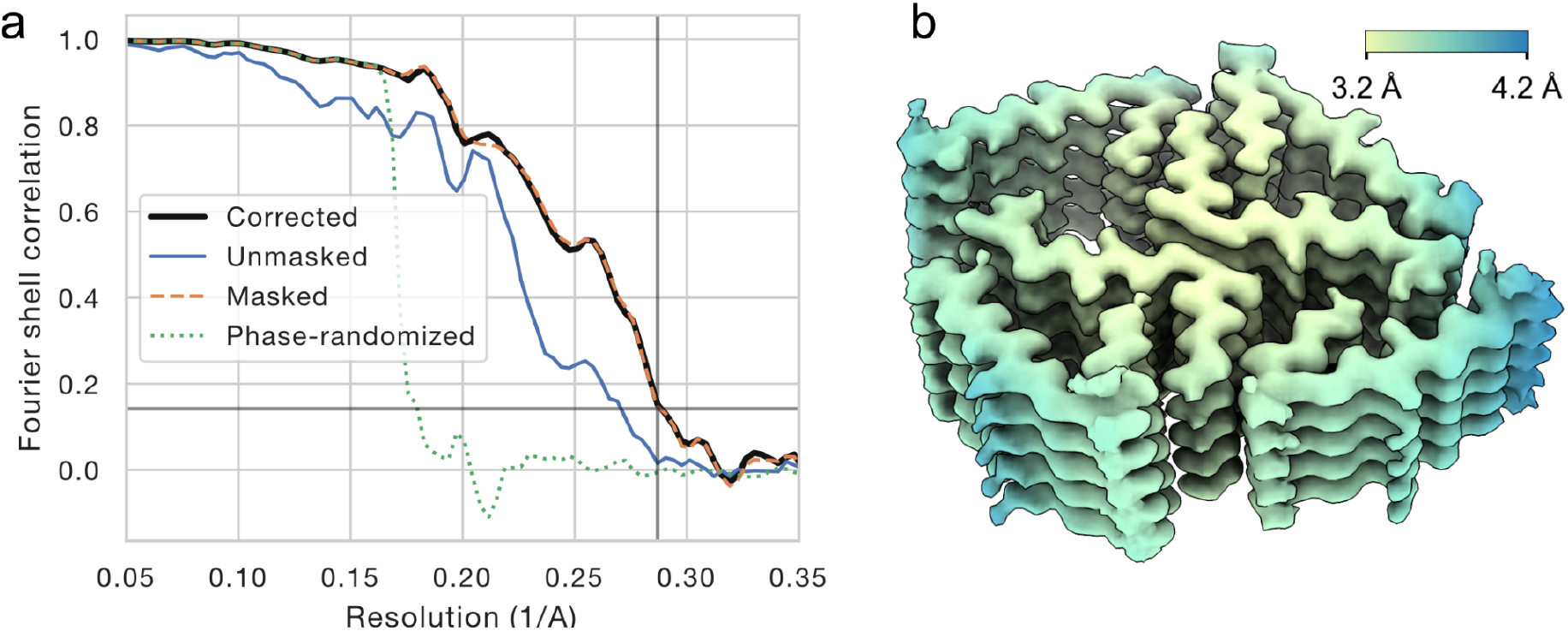
(a) Half-set Fourier shell correlation for aurein 3.3. The grey vertical line indicates the FSC=0.143 resolution of 3.48 Å, for a masked FSC after subtraction of a phase-randomized masked FSC (corrected, black line). (b) Local resolution map, calculated using half-set FSCs within a sliding window in Relion.

### Model Building

To build atomic models into the obtained maps, we proceeded identically for both uperin-3.5 type-I and aurein-3.3: Into each unique of the well-separated chains in the map, we built a peptide monomer manually using Coot 0.9.3^62^. While for aurein-3.3 all six unique chains are full-length, two out of the three unique chains in uperin-3.5 were truncated at their periphery after 14 residues (chain B) and 7 residues (chain C), respectively, as outside the tightly mated regions the map density became to blurry to trace the chains any further. We then expanded this set of monomers into three full helix layers in ChimeraX 1.3^63^, using the symmetry parameters determined in map refinement and the three-fold in-plane symmetry for the case of uperin-3.5. Interactive refinement guided by on-line molecular dynamics computations was then performed in Isolde 1.0b3^64^ within ChimeraX 1.2. Next, to mitigate edge effects of the molecular dynamics calculation, all helix layers but the central one were removed, and the single layer re-expanded to three layers, as above. This three-layer structure was then subjected to a final round of automatic real-space refinement in Phenix 1.19.1^65,66^, with imposing strict non-crystallographic symmetry restraints between equivalent chains. Model validation was carried out using MolProbity within Phenix 1.19.1^65,66^. The finally obtained model was further expanded to helices of arbitrary length using ChimeraX 1.3 for visualization, secondary structure analysis, and computation of SASA, coulomb, and lipophilicity potentials.

### Solvent-accessible surface area (SASA) calculations

The solvent-accessible surface area (SASA) buried per chain was calculated using the ‘measure buriedarea’ command in ChimeraX, and is defined by: ½ (sasa1 + sasa2 – sasa12) where sasa1 is the SASA enclosing the atoms in set1, sasa2 is the SASA enclosing the atoms of set2, and sasa12 is the SASA enclosing both sets of atoms together. The probe radius was 1.4 Å, often used to approximate a water molecule. Set1 was defined as a middle chain of a five-layered sheet, which is enough to completely cover the interactions of the middle chain. Set2 was defined as surrounding sheets as indicated in Table S1.

## Supplementary Information

**Supplementary Table 1:** Data collection parameters

|  | K3 dataset (Aurein) | K3 dataset (Uperin) |
| --- | --- | --- |
| Microscope | FEI Titan Krios G3i |  |
| Voltage (kV) | 300 |  |
| Grids | Type R2/1 Quantifoil |  |
| Magnification | npEFTEM SA105000 x | npEFTEM SA105000 x |
| C2 aperture size (μm) | 70 | 50 |
| Objective aperture size (μm) | No aperture | 100 |
| Pixel size (Å) | 0.85 Å/px |  |
| Total exposure (e-/Å<sup>2</sup>) | 40 |  |
| Exposure time (s) | 2 | 3 |
| Number of movie frames per exposure | 40 | 45 |
| Energy filter slit width (eV) | 15 | 20 |
| Data collection software | EPU 2.8.0.1256<br>Digital Micrograph 3.32.2403 |  |
| Number of exposures per hole | 5 |  |
| Defocus range (μm) | -0.5, -1.0, -1.5, -2, -2.5 | -1.0,-1.5,-2,-2.5,-3,-3.5,-4 |
| Number of micrographs collected/used for final map | 5514 / 482 | 11857 / 3297 |
| File Format | Tiff with LZW compression, gain corrected |  |

**Supplementary Table 2:** Software and algorithms used

| Name | Reference | Purpose |
| --- | --- | --- |
| MotionCor2 | https://emcore.ucsf.edu/ucsf-motioncor2/<br>(PMID:28250466)<sup>67</sup> | Motion correction of cryo-EM movies |
| crYOLO 1.8 | https://cryolo.readthedocs.io/<br>(PMID:32627734)<sup>59</sup> | Automatic filament picking in micrographs |
| Relion 3.1 | https://relion.readthedocs.io/<br>(PMID:32038040)<sup>68</sup> | Preprocessing, 2D and 3D classification, map auto-refinement, post-processing |
| CHEP | https://github.com/gschroe/chep<br>(PMID: 33556421)<sup>43</sup> | Filament polymorph classification |
| cryoSPARC 2.3 | https://cryosparc.com/<br>(PMID:28165473)<sup>60</sup> | Initial map building |
| Coot 0.93 | https://www2.mrc-lmb.cam.ac.uk/personal/pemsley/coot/<br>(PMID:20383002)<sup>62</sup> | Structure model building |
| UCSF ChimeraX 1.3 | https://www.rbvi.ucsf.edu/chimerax/<br>(PMID:28710774)<sup>69</sup> | Structure editing and visualization, model building |
| Isolde 1.0 | https://isolde.cimr.cam.ac.uk/<br>(PMID:29872003)<sup>64</sup> | Structure model building/refinement |
| Phenix 1.19.1 | https://www.phenix-online.org/<br>(PMID:20124702)<sup>65</sup> | Structure refinement and validation |

## Acknowledgments

This research was supported by the Ministry of Science, Research, Equalities and Districts of the Free and Hanseatic City of Hamburg (K.G., M.L., R.B.), Israel Science Foundation (grant no. 2111/20, M.L.), Israel Ministry of Science, Technology & Space (grant no. 3-15517, M.L.), U.S.-Israel Binational Science Foundation (BSF) (grant no. 2017280, M.L.), the Deutsche Forschungsgemeinschaft (DFG) grants INST 152/772-1,152/774-1, 152/775-1 and 152/776-1 FUGG to the CryoEM facility at CSSB and K.G., and the Joachim Herz Foundation (Add-on fellowship, R.B.). The work was performed at the Multi-User CryoEM Facility at the Centre for Structural Systems Biology, Hamburg. We thank the facility staff, especially Ulrike Laugks and Wolfgang Lugmayr, for their support.

## Notes

### Competing Interest Statement

The authors have declared no competing interest.

